# Rapid Restoration of Cell Phenotype and Matrix Forming Capacity Following Transient Nuclear Softening

**DOI:** 10.1101/2022.12.05.519160

**Authors:** Ryan C. Locke, Liane Miller, Elisabeth A. Lemmon, Sereen S. Assi, Dakota L. Jones, Eddie D. Bonnevie, Jason A. Burdick, Su Jin Heo, Robert L. Mauck

## Abstract

The dense extracellular matrix of connective tissues impedes cell migration and subsequent matrix formation at sites of injury. We recently employed transient nuclear softening via histone deacetylase inhibition with trichostatin A (TSA) treatment to overcome the stiff nuclear impediments to cell migration through dense tissues and electrospun matrices. Despite these positive findings, the long-term implications of transient nuclear softening on cell transcriptional phenotype and matrix formation capacity are unknown. To address this, we investigated the influence of transient TSA treatment on porcine meniscal cell behavior, beginning with the efficacy and reproducibility of transient TSA treatment on histone acetylation and chromatin remodeling in vitro and cell migration through native meniscus tissue. Within 3 days after cessation of transient TSA treatment, histone acetylation and chromatin remodeling returned to control levels. Following TSA treatment, endogenous cell migration through native meniscus tissue increased greater than 3-fold compared to controls. Importantly, meniscal cells completely restored their transcriptional phenotype and maintained their capacity to respond transcriptionally and functionally to a secondary pro-matrix stimuli (i.e., transforming growth factor β3) within 7 days after cessation of TSA treatment. Towards translation, we also showed the feasibility of biomaterial-delivered TSA to increase endogenous cell migration to a wound edge ex vivo. Together, this work defines the efficacy, reproducibility, safety, and feasibility of future translational approaches for nuclear softening to treat dense connective tissue injuries.

## Introduction

Following tissue injury, early colonization of the injured tissue site with cells that can deposit matrix is essential for endogenous tissue repair processes^1,2^. In soft tissues with loose collagen networks, wounds heal via a cascade of rapid inflammatory cell recruitment, endogenous interstitial (i.e., three-dimensional) cell migration and matrix production, wound edge contraction, and subsequent scar remodeling^3^. Conversely, in dense connective tissues, the high collagen network density impedes the critical step of interstitial cell migration to the injury site^3–5^. This results in reduced matrix production and limits wound healing outcomes in musculoskeletal tissues, such as the knee meniscus^6,7^ and articular cartilage^8,9^. Thus, therapeutic strategies that increase endogenous interstitial cell migration to the injury site have great potential to improve dense connective tissue repair.

Several strategies that either directly target cells or indirectly target the extracellular matrix (ECM) impediments to migration have been investigated to improve interstitial cell migration^10–12^. Beyond the introduction of migratory chemokine signaling gradients to induce cellular chemotaxis, we and others have developed therapeutic approaches that target mechanical impediments to migration of either the stiffness of 1) the ECM (therapeutic ‘microenvironment reprogramming’^10,11^) using degradative enzymes (e.g., collagenases) or 2) the cell nucleus (therapeutic ‘nuclear softening’^13–22^). In the former case, biomaterial-mediated delivery of collagenases increased the local ECM porosity, enabling enhanced interstitial cell migration.

Despite improvements, approaches that do not damage the native ECM are desirable. In the latter case, the nuclear elements that define stiffness, including the compaction state of the chromatin or intermediate filament molecules of the nuclear envelope, were targeted with histone deacetylase (HDAC) inhibitors or Lamin A/C knockdown, respectively^23–29^. The nuclear softening approach avoided damaging the native ECM, while also improving interstitial cell migration^13^.

We previously validated, both in vitro and in vivo, that nuclear softening via transient application of HDAC inhibitors (e.g., trichostatin A (TSA)) improved interstitial cell migration through dense musculoskeletal tissues and nanofibrous scaffolds^13^. However, despite the substantial benefits of nuclear softening for migration, this approach has the potential to irreversibly impact cellular phenotype^14,30,31^, negating the positive effects of the therapy. Ideally, nuclear softening would promote a transient increase in interstitial cell migration and then cells would recover their normal phenotype and regenerative capacity. Our recent studies suggest that this is the case, in that transient treatment (1 day) of the HDAC inhibitor TSA allowed for the recovery of normal histone acetylation levels within a few days^13^. However, it remains to be determined whether this acute HDAC inhibition irreversibly alters cell phenotype for regenerative matrix formation in dense connective tissue.

Here, in order to address this outstanding issue and begin translating these findings towards a clinically relevant in vivo model of meniscus repair in pigs, we assessed whether modulation of nuclear stiffness with TSA treatment in primary porcine meniscus cells resulted in chromatin remodeling, improved nuclear deformability, and enhanced migration through dense ECM. To assess the potential for phenotype recovery, we then evaluated the bulk transcriptional profile and matrix formation capacity of porcine meniscus cells with and without TSA treatment, including with subsequent treatment with a pro-matrix stimuli of transforming growth factor β3 (TGFβ3, **FIG. 1**). Finally, we evaluated the effect of localized biomaterial delivery of a nuclear softening agent (i.e., TSA) on the surrounding tissue and cells in a meniscus explant culture model. Overall, this work established transient nuclear softening of differentiated cells as a potential therapeutic avenue to dramatically increase interstitial migration of dense connective tissues cells without adversely impacting cellular phenotype or regenerative capacity for musculoskeletal tissue repair (**FIG. 1**).

**Figure 1.**
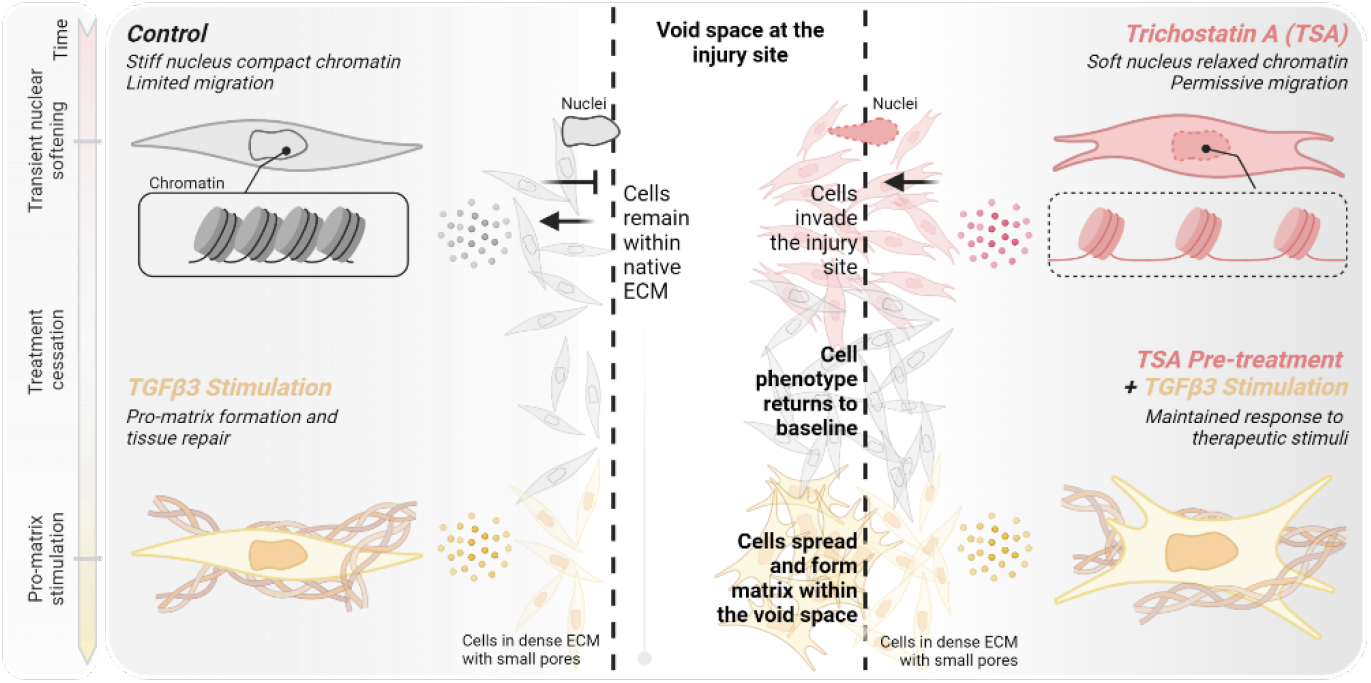
Study overview. We hypothesized that nuclear softening via transient trichostatin A (TSA) treatment promotes permissive cell migration into the void space at the injury site, while control cells remain within the injury site. Following cessation of TSA treatment, we hypothesized that cell phenotype returns to baseline and that cells pre-treated with TSA maintain the capacity to respond to subsequent therapeutic pro-matrix stimulation. Four illustrated treatment groups: 1) transient vehicle-treated control, 2) transient TSA, 3) transient vehicle pre-treatment plus subsequent transforming growth factor β3 (TBFβ3), and 4) transient TSA pre-treatment plus subsequent TGFβ3. Left timeline: the treatment course for transient nuclear softening via TSA, treatment cessation, and subsequent pro-matrix (e.g., TBFβ3) stimulation. Dashed lines: Small pores of dense extracellular matrix (ECM). Area between the dashed lines: the void space absent of reparative cells and matrix that forms following dense connective tissue injury. Flat arrowhead: inhibited interstitial migration through small pores. Callout boxes: Condensed and decondensed chromatin for control and TSA treated cells.

## Results

### Transient nuclear softening is effective and reproducible in porcine meniscus

In this study, as a step towards investigation in pre-clinical animal models in pigs, we sought to determine the 1) efficacy and reproducibility, 2) safety/mechanism (phenotype recovery), and 3) feasibility of nuclear softening to promote increased cell migration in porcine meniscus cells and tissue. To first establish the efficacy and reproducibility of TSA treatment, we probed the spatiotemporal effect of treatment with the histone deacetylase (HDAC) inhibitor, trichostatin A (TSA), on acetylation, chromatin condensation, nuclear deformability, and migration (2D and 3D) of primary porcine meniscus cells (**FIG. 2a**). For all of these outcome measures, cells were treated with medium-dose TSA at 325 nM for 24 hours based on our prior work in bovine meniscus cells^13^.

**Figure 2.**
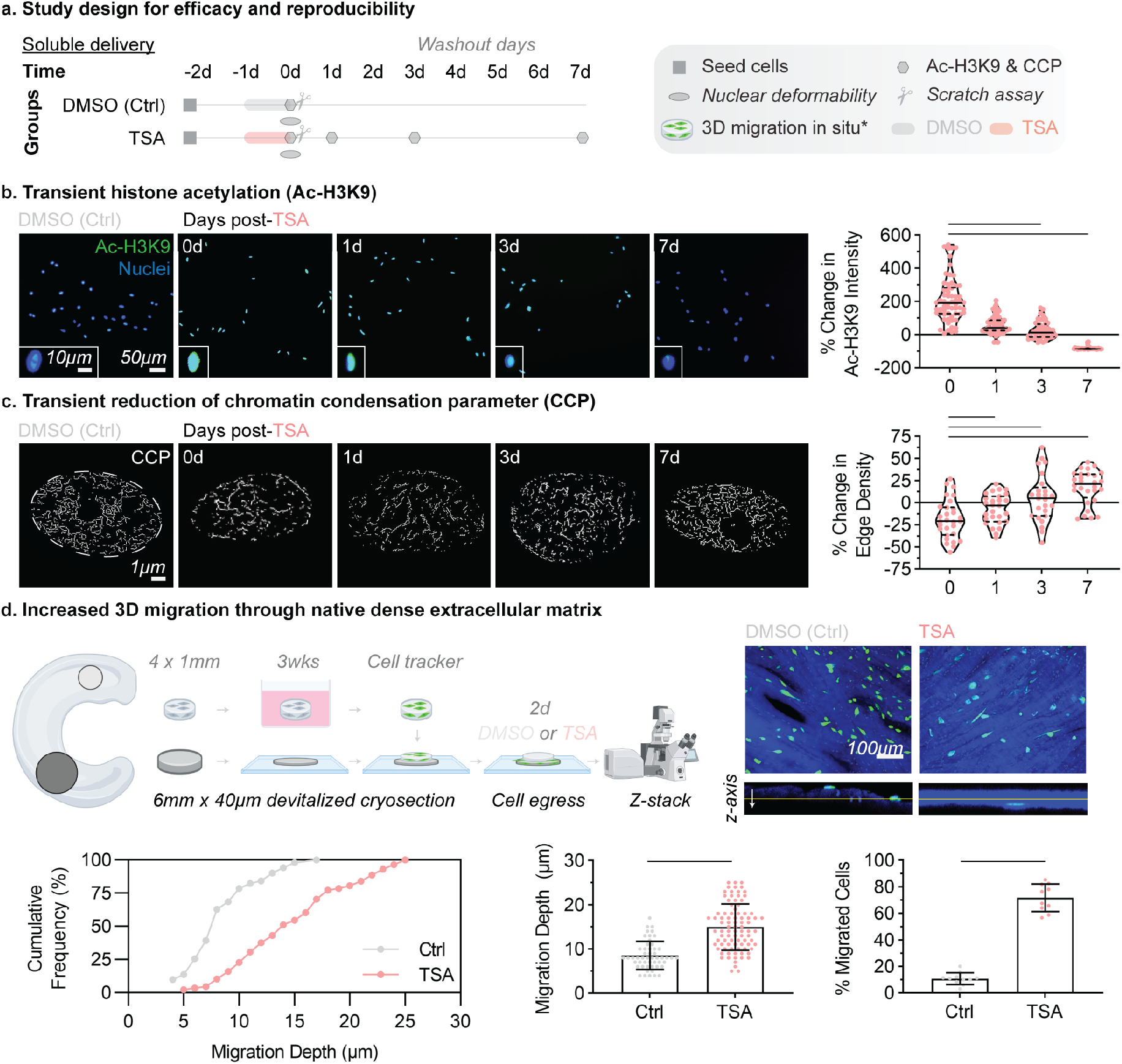
Nuclear softening is effective and reproducible. (a) Study design to confirm the efficacy and reproducibility of nuclear softening in terms of increased histone acetylation, chromatin decondensation, nuclear deformability, and migration in porcine meniscus cells. TSA: trichostatin A, DMSO: dimethyl sulfoxide, TGFβ3: transforming growth factor β3, Ctrl: Control. Highlighted lines: treatments of DMSO (gray), TSA (red), and TGFβ3 (yellow). Shapes: specific start and end points denoted in the legend. d: day. Ac-H3K9: acetylated H3K9, CCP: chromatin condensation parameter. (b) Histone acetylation in DMSO-treated controls and 0, 1, 3, and 7 days post-TSA treatment. Fluorescence images showing Ac-H3K9 (green) and DAPI counterstain (blue). Scale bar = 50µm. Inset images: Zoom-ins of representative nuclei at each time point. Inset scale bar = 10µm. Quantification of percent change in Ac-H3K9 intensity at each time point following TSA treatment normalized to DMSO-treated controls. 0% change: identical values to baseline DMSO-treated control. TSA: trichostatin A, DMSO: dimethyl sulfoxide. Bars: significantly different, p<0.05. One-way ANOVA. (c) Transient chromatin remodeling and relaxation in DMSO-treated controls and 0, 1, 3, and 7 days post-TSA treatment. Edge detection from high magnification imaging of DAPI-stained nuclei. Scale bar = 1µm. Dotted circle: boundary of DAPI-stained nuclei. Quantification of percent change in chromatin condensation parameter (CCP) normalized to DMSO-treated controls. Bars: significantly different, p<0.05. One-way ANOVA. (d) Nuclear softening with TSA enhances interstitial migration of MFCs through native tissues. Schematic showing native meniscal tissue explants and devitalized tissue sections for 3D migration assays. Cell egress from the explant and infiltration into the devitalized tissue section was evaluated by z-stacks from confocal microscopy. Max intensity projection images and cross-sectional views of DMSO-treated control and TSA-treated constructs showing migrating pMFCs (green) and the devitalized tissue sections (blue). Quantification of cumulative frequency, migration depth, and percent migrated cells with or without TSA treatment. Bars: significantly different, p<0.05. Unpaired t-test.

we first quantified the acetylation level of lysine 9 on histone 3 (Ac-H3K9) in cells treated with TSA or untreated controls. Following treatment with TSA for 24 hours (day 1), Ac-H3K9 significantly increased compared to controls (**FIG. 2b**). Following removal of TSA, cells were further cultured in basal media for up to 7 days (**FIG. 2b**). To assess persistence of treatment, Ac-H3K9 was quantified at intervals of 1, 3, or 7 days following removal of TSA (**FIG. 2b**). Consistent with our previous findings, Ac-H3K9 remained increased 1 day after treatment compared to controls, and slowly decreased to control levels by 3 days after TSA removal (**FIG. 2b**).

To evaluate the change in chromatin condensation with TSA treatment, we imaged individual cell nuclei at high resolution by confocal microscopy and calculated the chromatin condensation parameter (CCP) based on edge detection within 4’,6-diamidino-2-phenylindole (DAPI) stained nuclei^32^. Following treatment with TSA for 24 hours, CCP decreased by 20% compared to control cells (**FIG. 2c, SUPP. FIG. 1a**). Following removal of TSA, CCP increased to control levels within 3 days, indicating reestablishment of homeostatic chromatin organization from the previously relaxed state (**FIG. 2c, SUPP. FIG. 1a**). To assess whether the changes in chromatin condensation from a more condensed to a more relaxed state altered the nuclear response to applied deformations (i.e., the nuclear deformability), cells were seeded on nanofibrous scaffolds for 24 hours then treated with TSA for 24 hours. After TSA treatment, cell-seeded scaffolds were stretched from 0 to 15% grip-to-grip strain and the change in nuclear aspect ratio (NAR) was quantified (**SUPP. FIG. 1b**). NAR correlated with increasing strain for scaffolds treated with TSA compared to control scaffolds (**SUPP. FIG. 1b**), indicating increased nuclear deformability after TSA treatment.

To evaluate the effect of chromatin remodeling and nuclear deformability on cell migration, we assessed 2-dimensional migration using the scratch assay and 3-dimensional migration of tissue resident cells using explanted meniscus biopsies. Following monolayer scratch in vitro, cells treated with TSA showed no difference in scratch closure rate compared to controls (**SUPP. FIG. 1c**). We next evaluated meniscus cell migration through the dense fibrous ECM of meniscus tissue. To do so, porcine meniscus explants were cultured for 3 weeks, and then endogenous cells were stained with a fluorescent dye to track cells over culture time (**FIG. 2d**). To investigate cell egress from the explant and migration, explants were set on top of flat sections of devitalized meniscus tissue (termed explant-ECM samples). Explant-ECM samples were cultured for 2 days with or without continuous TSA (**FIG. 2d**), and migrated cells within the devitalized explant were imaged using confocal microscopy. Without TSA, cells migrated out of the explants but were located predominantly at the tissue surface (**FIG. 2d**). Conversely, with TSA treatment, cells migrated out of the explants and into the ECM (**FIG. 2d**). The cumulative frequency, migration depth, and percent migrated cells into the devitalized meniscus ECM significantly increased with TSA treatment compared to controls (**FIG. 2d**). Together, these results establish the efficacy and reproducibility of transient nuclear softening in increasing interstitial cell migration through dense ECM.

### Following transient nuclear softening, cells restored their native transcriptional profile

While the above data indicated that TSA promotes cell migration, the phenotypic status of cells post-TSA exposure remains an open question. Therefore, to determine the safety/mechanism of nuclear softening, we queried cellular phenotype by carrying out bulk transcriptomic analysis and assessing matrix formation capacity of porcine meniscus cells after TSA treatment. For RNA-sequencing and nascent matrix formation, we fabricated aligned nanofibrous polycaprolactone scaffolds via electrospinning and coated the scaffolds in fibronectin to promote cell attachment. We isolated meniscus cells from 3 porcine donors (N=3, 6-9 months old) and seeded them onto scaffolds for 24 hours prior to TSA treatment (day -2, **FIG. 3a**). At 24 hours (day -1, **FIG. 3a**), scaffolds were treated with TSA at either low dose (150 nM) or high dose (650 nM) for an additional 24 hours. The low dose was chosen based on our prior work, which showed that 150 nM was the lowest TSA dose to increase 3-dimensional cell migration^13^. The high dose was chosen based on prior work using 650 nM, as well as evidence that >1 uM TSA treatment reduced metabolic activity both in vitro and ex vivo^13^. Following removal of TSA (day 0), cells were cultured in basal media for up to 7 days (**FIG. 3a**).

**Figure 3.**
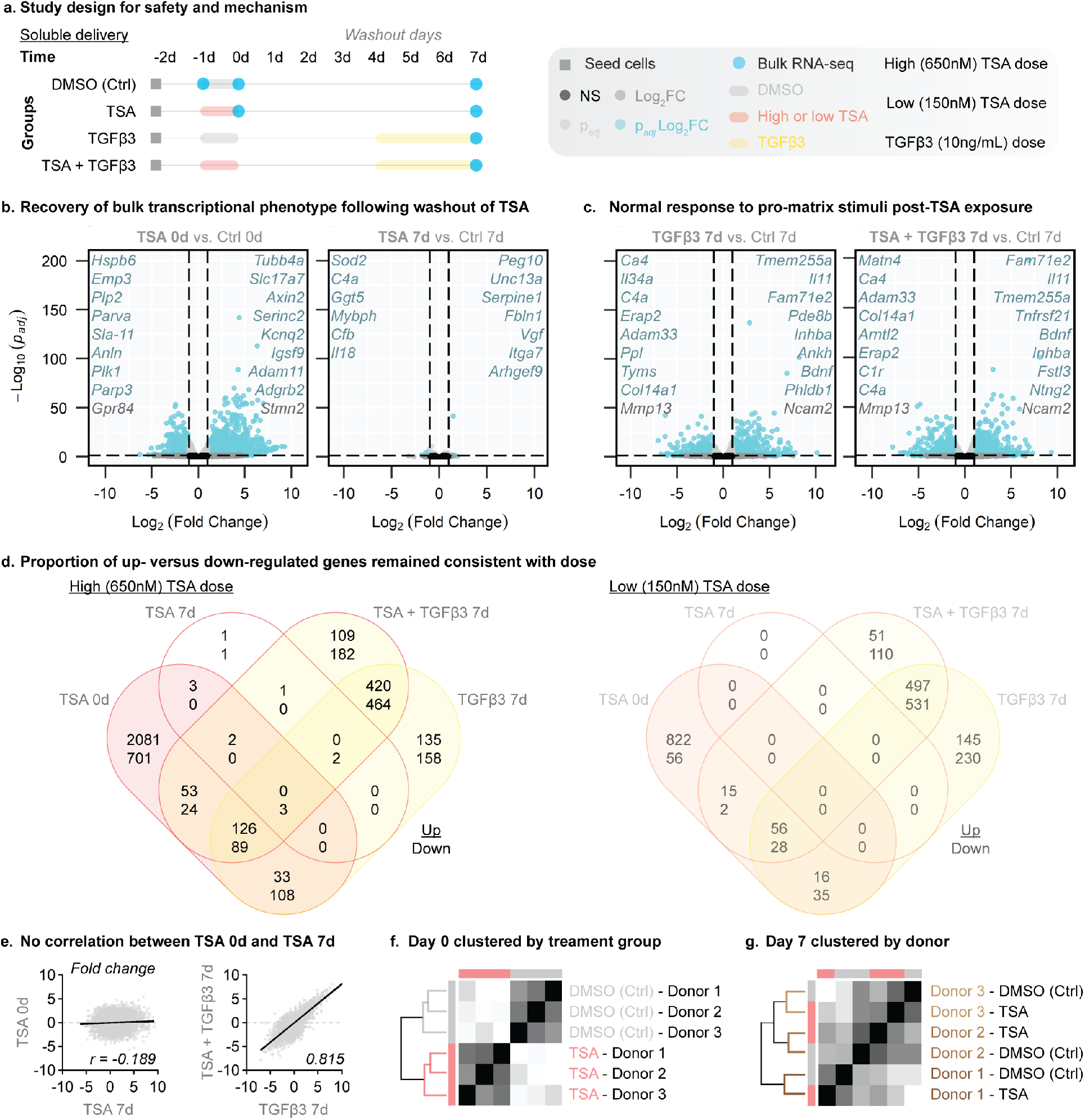
RNA-sequencing following transient nuclear softening demonstrates recovery in transcriptional profile. (a) Study design to evaluate safety and mechanism of nuclear softening on porcine meniscus cells cultured on fibrous scaffolds using bulk RNA-sequencing (RNA-seq). TSA: trichostatin A, DMSO: dimethyl sulfoxide, TGFβ3: transforming growth factor β3, Ctrl: Control. Highlighted lines: treatments of DMSO (gray), TSA (red), and TGFβ3 (yellow). Shapes: specific start and end points denoted in the legend. d: day. (b-c) Volcano plots showing differentially expressed genes between treatment groups (blue dots at 0d or 7d). Vertical dashed line: p_adj_<0.05, horizontal dashed lines: -1 > Log_2_(FC: Fold Change) > 1, NS: not significant. (b) Volcano plots of 0 days post-TSA compared to time-matched 0 day DMSO-treated control (Ctrl) and of 7 day post-TSA compared to time-matched 7 day DMSO-treated control. (c) Volcano plots of TGFβ3 treatment at 7 days compared to time-matched 7 day DMSO-treated control (Ctrl) and of 7 day post-TSA + TGFβ3 compared to time-matched 7 day DMSO-treated control. Highlighted lines: duration of treatments with DMSO (gray), TSA (red), and TGFβ3 (yellow). d: day. Light blue genes: differentially expressed genes with p_adj_<0.05 and -2 > fold change > 2. Italicized gene names: top up- and down-regulated genes for each group. (d) Venn diagrams of differentially expressed genes between groups for high-dose and low-dose TSA treatments. The number of upregulated (top number) and downregulated (bottom number) genes. Non-overlapping quadrants represent unique gene sets for each group. (e) Correlation of fold changes between TSA 0d and TSA 7d and between TSA + TGFβ3 7d versus TGFβ3 7d. (f-g) Expression patterns were hierarchically clustered by (f) treatment group at immediately following TSA treatment and (g) by donor at day 7 post-TSA.

In addition to treatment with TSA, a group of scaffolds were also continuously treated with TGFβ3 at day 4 until day 7 (**FIG. 3a**). The endpoint was selected based on our previous data in bovine cells that showed recovery of baseline acetylation between 5 and 7 days after TSA treatment^13^. RNA-sequencing was performed on days -1 (before TSA treatment), 0 (immediately after TSA treatment), and on day 7 (**FIG. 3a**). Nascent matrix production was assessed at day 7 and at day 14 to assess longer term function (**FIG. 5a, SUPP. FIG. 2a**).

With either a low- or high-dose TSA treatment, doses that promoted increased cell migration in our prior work^13^, we identified both time- and dose-dependent effects on meniscus cell phenotype (**FIG. 3b, SUPP. FIG. 3c-d**). Strikingly, cells treated with low-dose TSA for 24 hours completely restored their native phenotype based on the complete absence of differentially expressed genes within 7 days following removal of TSA (**SUPP. FIG. 3a**). Likewise, cells treated with high-dose TSA for 24 hours nearly restored their native phenotype within 7 days after TSA treatment, with only 13 differentially expressed genes remaining (**FIG. 3b**). Therefore, we focused on the high-dose TSA treatment to better assess the safety and any potential long-term effects of TSA treatment.

Immediately after high-dose TSA treatment (day 0), 2,298 genes were upregulated, and 925 genes were downregulated, compared to time-matched controls (**FIG. 3b**). Notably, all but 13 genes returned to time-matched control levels after high-dose TSA removal and 6 additional days in basal media (**FIG. 3b**). Importantly, cells treated with TSA also maintained a normal transcriptional response to the pro-matrix stimuli of TGFβ3 (**FIG. 3c**). In particular, the top up- (e.g., *Ncam2, Il11*) and down- (e.g., *Mmp13, Adam33*) regulated genes overlapped substantially after TGFβ3 treatment whether alone or after TSA + TGFβ3 treatment (**FIG. 3c**).

To identify unique and overlapping genes between the time points and treatment groups, we compared the differentially expressed genes (i.e., genes with adjusted p-value < 0.05 and fold change > 2) between all groups for low- and high-dose TSA (**FIG. 3d**) and between low- and high-dose for matching groups (e.g., low-dose TSA 0d vs. high-dose TSA 0d, **SUPP. FIG. 3b-d**). For low- and high-dose TSA, the proportion of up- and down-regulated genes remained consistent for all groups, with less differentially expressed genes following low-dose compared to high-dose TSA, as expected (**FIG. 3d**). Of the 1,030 differentially expressed genes at day 0 following low-dose TSA, 91% were also differentially expressed at day 0 following high-dose TSA (**SUPP. FIG. 3b**). This resulted in 97 (83 up- and 14 down-regulated) and 2,290 (1,142 up- and 818 down-regulated) unique differentially expressed genes at day 0 for low-dose and high-dose TSA treatment, respectively (**SUPP. FIG. 3b**). Notably, only 2 of the 13 differentially expressed genes at day 7 after high-dose TSA removal were unique to TSA treatment (**FIG. 3d**), indicating minimal lasting effect of prior TSA exposure on cell phenotype.

To compare the gene expression relationship more broadly over time and following induction of matrix formation, we evaluated the correlation (**FIG. 3e**) and unbiased hierarchical clustering based on treatment group (TSA vs. DMSO) and porcine donor (**FIG. 3f-g**). Correlation plots comparing all differentially regulated genes showed a significant negative correlation at 0 compared to 7 days after high-dose TSA treatment (Pearson’s r = -0.189, **FIG. 3e**). Conversely, a strong positive correlation (Pearson’s r = 0.815) was observed for differentially regulated genes in the high-dose TSA + TGFβ3 treatment compared to TGFβ3 treatment alone groups (**FIG. 3e**). These correlation data indicate no similarity in phenotype between 0 and 7 days following high-dose TSA treatment, and that prior TSA exposure does not perturb the cellular response to pro-matrix agents, such as TGFβ3 (**FIG. 3e**). From hierarchical clustering of gene expression patterns across treatment groups and donors, we identified that gene expression clustered by treatment group immediately after high-dose TSA on day 0 (**FIG. 3f**) and by donor after washout of TSA on day 7 (**FIG. 3g**). This further indicates that at 7 days following TSA exposure the cell phenotype mirrors that of healthy vehicle-treated control cells.

### Nuclear softening did not permanently alter phenotypic gene pathways

From the differentially expressed genes from each group (**FIG. 3c**), we evaluated the significantly enriched biological processes using the DAVID Functional Annotation Tool (**FIG. 4**)^33–37^. For all genes immediately downregulated following high-dose TSA treatment, biological processes related to the cell cycle, chromosomes, and DNA were enriched (**FIG. 4a**). For all genes immediately upregulated following high-dose TSA treatment, ontologies related to ion transport were enriched, with regulation of ion transmembrane transport being the most enriched ontology (**FIG. 4a’**). When evaluated at 7 days following removal of the transient TSA treatment, no ontologies were enriched for the upregulated genes, and inflammatory response was the only enriched ontology for the downregulated genes (**FIG. 4b, b’**). Importantly, the majority of differences in the gene expression profile at day 0 were absent at day 7 following TSA, suggesting a near complete reversal back to naïve cell phenotype (**FIG. 4e-f**). These data support that transient treatment with TSA has no lasting effect on cell phenotype in the long term.

**Figure 4.**
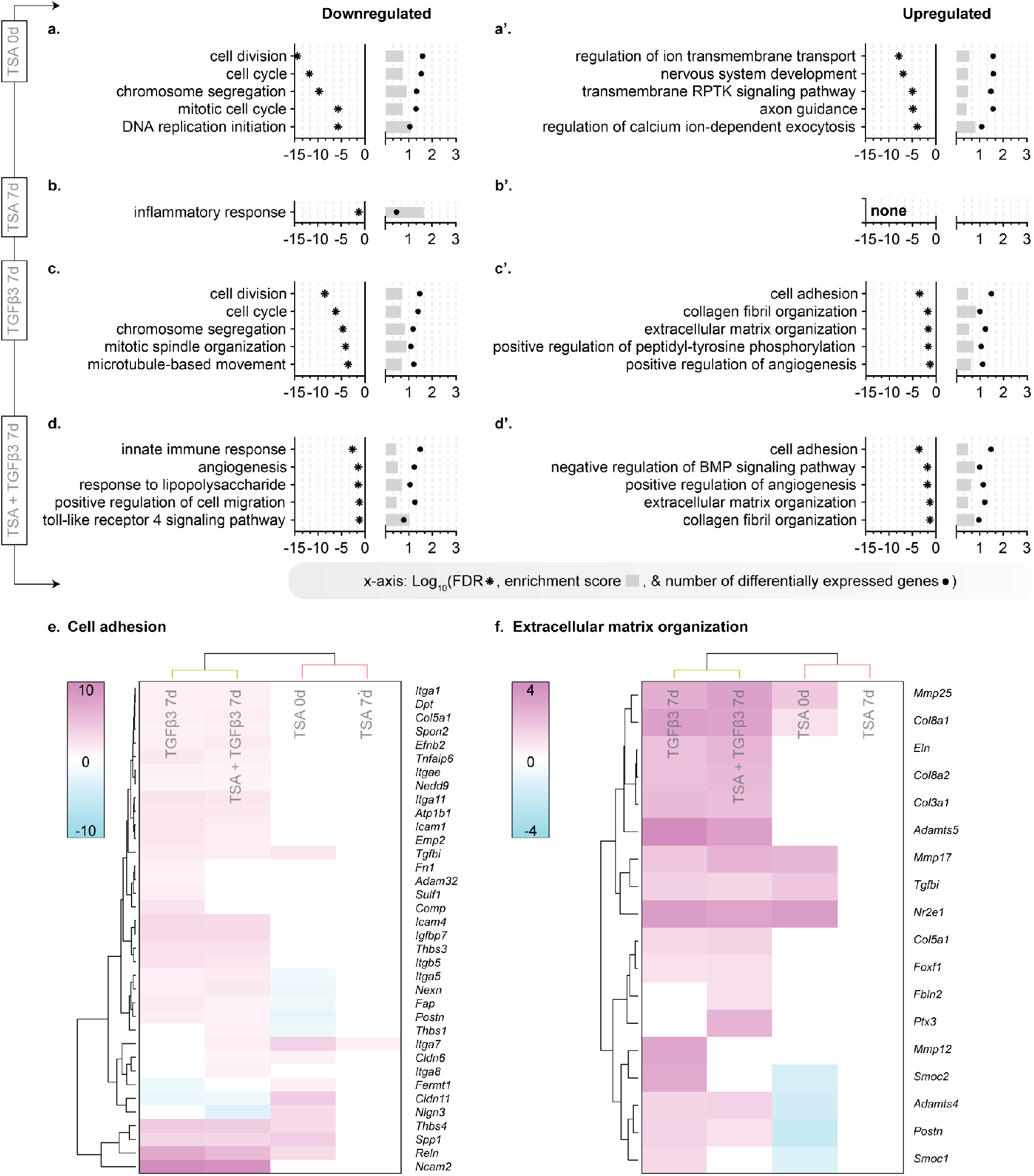
Nuclear softening did not permanently alter phenotypic gene pathways. Gene ontologies for down- (a, b, c, d) and up- (a’, b’, c’, d’) regulated differentially expressed genes in each group. Note: No gene ontologies were significantly enriched for the upregulated genes from the TSA 7d group. DNA: deoxyribonucleic acid, RPTK: receptor protein tyrosine kinase, BMP: bone morphogenic protein. Heatmaps for the enriched ontologies of (e) cell adhesion and (f) extracellular matrix organization with hierarchical clustering of treatment group (columns) and genes (rows). Yellow bar: Clustered by TGFβ3. Light red bar: Clustered by TSA. Heatmap scales: Fold change in gene expression compared to time-matched control with 0 representing genes that were not differentially expressed. Significance: FDR<0.05, from DAVID analysis.

**Figure 5.**
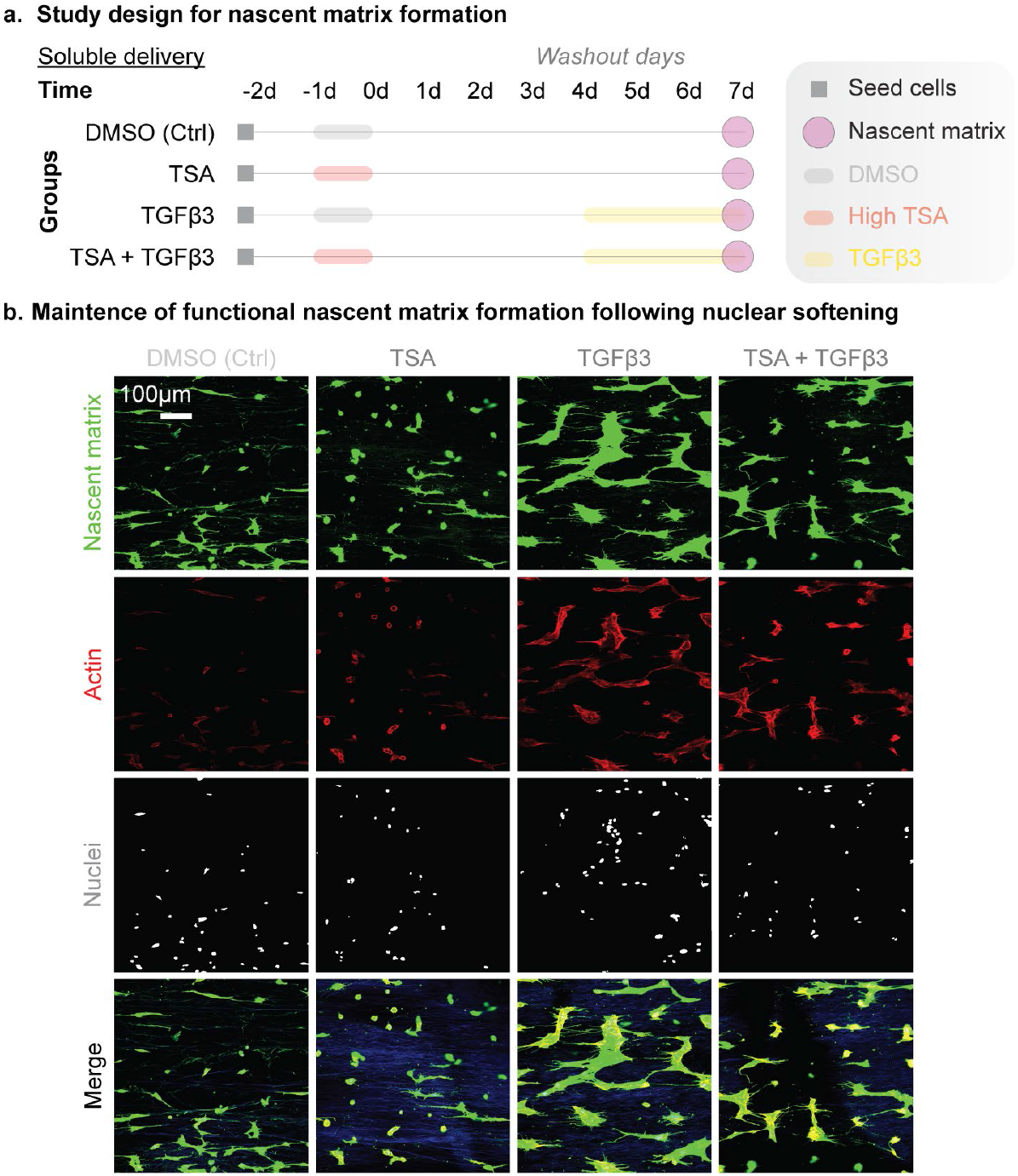
Following transient nuclear softening, meniscus cells maintained matrix formation capacity. (a) Study design to evaluate the functional effect of nuclear softening on nascent matrix production in porcine meniscus cells cultured on fibrous scaffolds. TSA: trichostatin A, DMSO: dimethyl sulfoxide, TGFβ3: transforming growth factor β3, Ctrl: Control. Highlighted lines: treatments of DMSO (gray), TSA (red), and TGFβ3 (yellow). Shapes: specific start and end points denoted in the legend. d: day. (b) Representative confocal images of nascent matrix at 7 days following DMSO, TSA, TGFβ3, or TSA + TGFβ3 treatment. Green: deposited nascent matrix over the culture period, yellow: overlay of nascent matrix and actin (red), blue: nanofibrous scaffold autofluorescence, magenta: nuclei.

We further probed whether cells could respond to anabolic stimulation following TSA treatment using this same approach. The enriched ontologies for down- (**FIG. 4c, d**) and up- (**FIG. 4c’, d’**) regulated genes following TGFβ3 treatment (**FIG. 4c, c’**) or high-dose TSA + TGFβ3 treatment (**FIG. 4d, d’**) on day 7 were comparable to one another. The enriched ontologies for high-dose TSA + TGFβ3 treatment closely mirrored the enriched ontologies for TGFβ3 treatment alone, suggesting prior TSA treatment does not alter the cellular response to stimuli that promote matrix formation. In fact, the gene expression profiles for high-dose TSA + TGFβ3 and TGFβ3 treatment alone clustered together for gene groups related to cell adhesion (**FIG. 4e**) and extracellular matrix organization (**FIG. 4f**), indicating a similar transcriptional response to TGFβ3 treatment in both groups. This is evident in the substantial overlap seen in differential expression for all groups treated with TGFβ3 (**SUPP. FIG. 4a**). Although all of the groups treated with TGFβ3 did not identically overlap in differential expression (**SUPP. FIG. 4a**), only the biological processes for the unique downregulated genes following TGFβ3 treatment were significantly different (**SUPP. FIG. 4b**). No biological processes were significantly different for both the unique up- and down regulated genes following high-dose TSA + TGFβ3 or low-dose TSA + TGFβ3 (**SUPP. FIG. 4a-b**). While beyond the scope of this study, the differences between the downregulated genes following TGFβ3 treatment only compared to the groups with prior TSA exposure remains an area for future investigations.

### Cells retain the ability to produce nascent matrix after transient nuclear softening

In addition to cell phenotype, we assessed whether there were functional consequences of high-dose TSA treatment by evaluating matrix formation capacity, which is necessary for wound repair (**FIG. 5ac**). Using the same treatment regimen as above, we labeled the nascent matrix formed by cells on electrospun scaffolds. Cell-seeded scaffolds were cultured in defined media that included the noncanonical amino acid azidohomoalanine (AHA) and methionine at a ratio 3:1, respectively. During the culture, cells incorporate the azide-modified noncanonical amino acid into any new cell-produced protein (i.e., the nascent matrix) that requires methionine during synthesis, enabling their subsequent identification via a click chemistry mediated azide-alkyne cycloaddition with an alkyne-containing dye^38–41^. Results from this analysis at day 7 showed that cells treated with high-dose TSA produced similar levels of nascent matrix compared to DMSO-treated controls (**FIG. 5b**) and were equally responsive to TGFβ3 stimulation (**FIG. 5b**), similar to prior work in human ventricular fibroblasts^14^. At day 14, these findings persisted, with all groups increasing in the amount of labeled nascent matrix compared to day 7 (**SUPP. FIG. 2**). These results indicate that transient nuclear softening does not damage the cells’ ability to produce matrix for tissue repair in the longer term.

### Localized and rapid delivery of nuclear softening agent using dual-material scaffold increased acetylation and cellularity at the wound edge

Lastly, as a proof-of-concept, and to show the feasibility of locally delivering nuclear softening agents to meniscus injury sites, we fabricated dual electrospun fibrous scaffolds^13^ that support cell attachment as well as rapidly release TSA (**FIG. 6a**). To fabricate the scaffolds, we dual electrospun polycaprolactone (PCL, long-term stable fiber fraction) and TSA loaded into water soluble poly(ethylene oxide) (PEO, sacrificial fiber fraction for increased porosity and payload delivery). Upon exposure to aqueous environments, the PEO rapidly dissolves to release the TSA locally at the delivery site (**FIG. 6a**). The composite scaffolds were delivered into vertical defects in porcine meniscus explants, sutured in place, and cultured for 1 or 7 days in basal media (**FIG. 6a**). Following culture, we quantified the local acetylation and cellularity at the defect edges to assess therapeutic efficacy.

**Figure 6.**
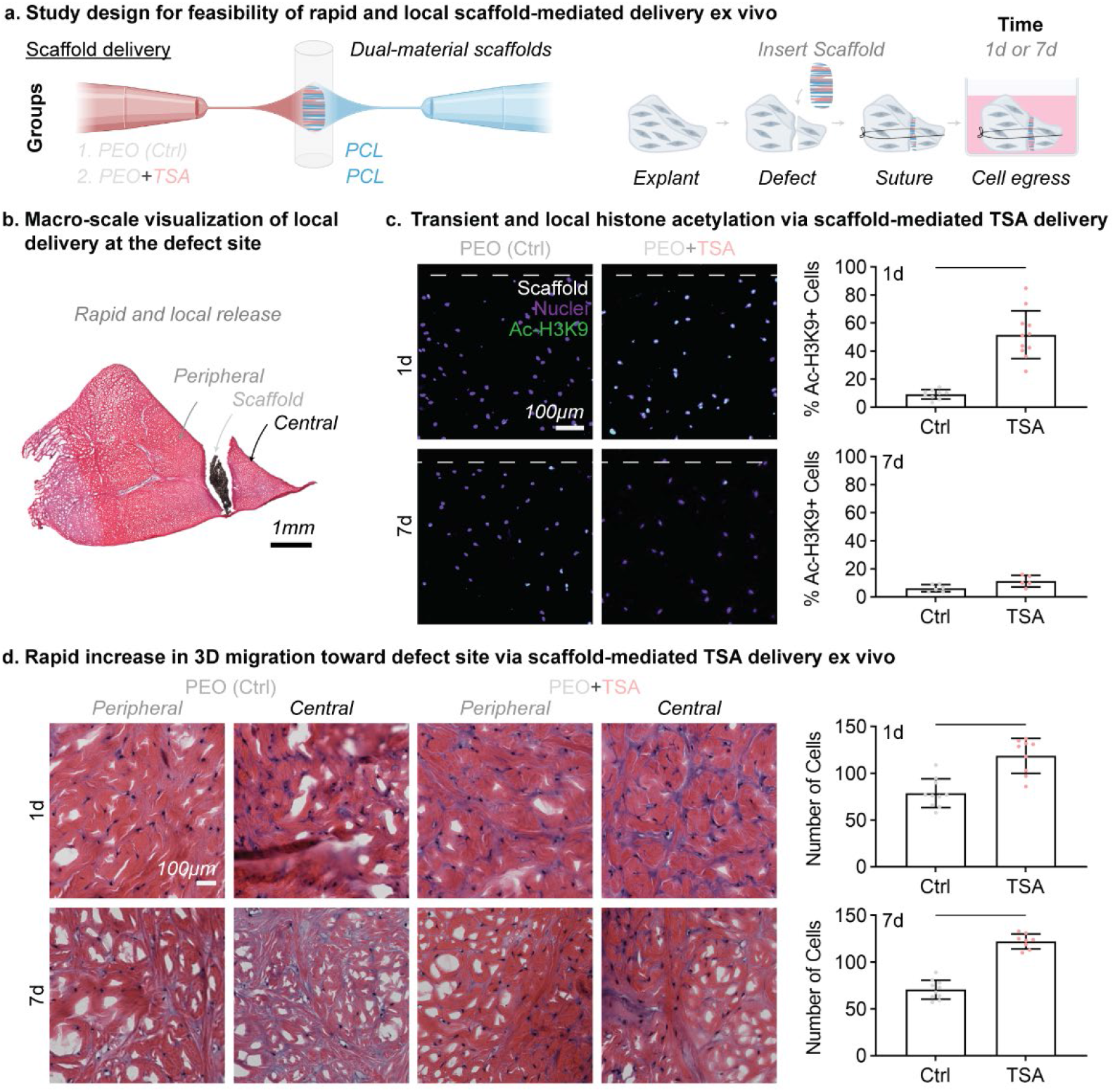
Scaffold-mediated TSA delivery rapidly and locally increased cell migration to the meniscal wound edge in an ex vivo injury model. (a) Study design to test the feasibility of nuclear softening using a dual-material scaffold loaded with TSA for local delivery to the wound site in porcine meniscus explants. PCL: polycaprolactone, PEO: polyethylene oxide. d: day. (b) Schematic showing dual-material polycaprolactone (PCL) and polyethylene oxide (PEO) nanofibrous scaffold fabrication and delivery. Representative cross section of vertical defects in whole meniscal explants and localization of the dual-material scaffolds within the defects and sutured in place, showing the peripheral and central margins (c) Fluorescent images of acetylated H3K9 (Ac-H3K9) in explant cryosections. Dashed line: scaffold location. Quantification of the percent of Ac-H3K9 positive cells following 1 and 7 days of culture for explants treated with PEO-control or TSA-loaded scaffolds. Scale bar = 100µm. Bar: significant difference, p<0.05. Unpaired t-test. (d) Hematoxylin (magenta: nuclei) and eosin (pink: extracellular matrix) stained sections following 1 and 7 days of culture for explants treated with PEO-control or TSA-loaded scaffolds. Total number of cells (peripheral and central wound edges) were quantified at 1 and 7 days of culture for each group. Bars: significant difference, p<0.05. Unpaired t-test.

To determine the localization of outcomes related to scaffold-delivered TSA, a vertical meniscus defect was created in the porcine meniscus and scaffolds either with (PEO+TSA) or without TSA (PEO only control) were sutured into the defect (**FIG. 6a**). These constructs were cultured for either 1 day to assess the immediate response to TSA or for 7 days to assess the persistence of TSA treatment after delivery (**FIG. 6a**). The constructs were cryosectioned, stained, and imaged at the peripheral or central margins (**FIG. 6b**). Immunofluorescent imaging of Ac-H3K9 in meniscus cryosections showed that local TSA delivery resulted in higher levels of Ac-H3K9 compared to controls after 1 day (**FIG. 6c**). By day 7, acetylation levels returned to control levels (**FIG. 6c**), suggesting successful transient local delivery of TSA. From histological sections stained with hematoxylin and eosin, we also noted increased cellularity at the wound interface for TSA-treated constructs compared to controls after only 1 day (**FIG. 6d**). This rapid and local increase in cellularity was maintained over 7 days (**FIG. 6d**). With feasibility to rapidly increase cellularity in a clinically relevant ex vivo model, this biomaterial delivery system motivates the continued investigation of therapeutic transient nuclear softening for in vivo tissue repair.

## Discussion

This study validated that transient chromatin decondensation using the histone deacetylase (HDAC) inhibitor trichostatin A (TSA) does not cause irreversible changes in meniscus cell acetylation, transcriptional profile, or matrix production capacity. Taken together, these data supports that nuclear softening is not only effective at improving cell migration, but also a safe and practical approach to improving wound site cellularity.

Specifically, we first confirmed the efficacy and reproducibility of chromatin decondensation to ‘soften’ the nucleus as well as improve interstitial migration capacity in the dense three-dimensional matrix of native meniscal tissue. Since our initial work was in less translatable bovine meniscus cells, here we used translatable porcine meniscus cells and showed that these cells mirrored the prior results in terms of increased histone acetylation, chromatin decondensation, and 3D interstitial cell migration following TSA treatment. Additionally, these TSA-mediated alterations in chromatin condensation and histone acetylation were transient, not permanent, with cells returning to control levels within 7 days of treatment cessation. Moreover, nuclear softening enhanced endogenous 3D migration through their native dense meniscus ECM, indicating the therapeutic potential of nuclear softening to improve the repair capacity of dense musculoskeletal tissues through enhanced interstitial cell migration to the wound edge. From bulk RNA-sequencing, the immediate transcriptional response following nuclear softening was reversed with complete transcriptional recovery by 7 days, highlighting the safety of low-dose nuclear softening. We further identified that prior TSA treatment does not significantly alter the transcriptional or functional response to subsequent pro-matrix stimuli. Thus, future work may safely explore combinatorial approaches using primary nuclear softening enhanced by a secondary therapeutic stimuli to improve matrix formation and repair outcomes. Lastly, to further establish the translational feasibility of nuclear softening, we showed that local, biomaterial-mediated delivery of TSA resulted in rapid and transient increases in acetylation and interstitial cell migration in an in vitro meniscus repair model, with effects localized to the wound edges.

## Conclusions

This work shows the transcriptional recovery of primary meniscus cells following transient histone deacetylase inhibition. Small molecules that increase cell migration while preserving matrix production capacity without aberrant or prolonged changes to native cell phenotype hold promise for regenerative treatments for meniscus injury and across multiple dense connective tissues.

## Methods

### Primary cell isolation and culture

Porcine menisci (6-8 months old) were dissected from cadavers within four hours of euthanasia and incubated in a basal media consisting of Dulbecco’s modified Eagle’s medium (DMEM) with 10% fetal bovine serum (FBS) and 1% penicillin/streptomycin/fungizone (PSF). Primary porcine meniscus fibrochondrocytes (pMFC) were isolated by first mincing menisci into ∼1 mm segments followed by culture on tissue culture plastic and incubation at 37 °C in basal media with media changes every other day. Once egressed cells reached 70-80% confluency, passage 0 cells were harvested using 0.25% trypsin and then expanded or frozen. Passage 1-2 cells were used for all experiments.

### Aligned nanofibrous PCL scaffolds

Aligned nanofibrous poly(ε-caprolactone) (PCL) scaffolds were fabricated via electrospinning. Briefly, a PCL solution (80 kDa, Shenzhen Bright China Industrial Co., Ltd., China, 14.3% wt/vol in a 1:1 solution of tetrahydrofuran and N,N-dimethylformamide) was extruded (2.5 mL/hour) through a stainless steel 18G needle t charged to +15 kV. To form aligned scaffolds, fibers were collected on a mandrel rotating with a high surface velocity (∼10 m/s). The scaffolds were hydrated and sterilized in ethanol in a stepwise fashion starting with 100%, then 70%, 50%, and 30% ethanol for 30 minutes, followed by two washes in sterile PBS with 1% PSF prior to cell seeding.

### Aligned nanofibrous dual PCL/PEO scaffolds

Using co-electrospinning, aligned nanofibrous dual-component scaffolds were developed with approximately a 50:50 ratio of PCL and water-soluble polyethylene oxide (PEO; 200 kDa, Polysciences, Inc., Warrington, PA). Briefly, PCL (formulated as described above) and PEO (10% wt/vol in 90% ethanol) solutions were electrospun simultaneously onto a centrally placed mandrel (using the same electrospinning parameters described above). To fabricate scaffolds containing the histone deacetylase inhibitor (trichostatin A, TSA, Catalog #: T8552-1MG, Sigma Aldrich), the PEO solution was supplemented with 1 % wt/vol of TSA. To develop PCL/PEO-TSA scaffolds, PCL (10 mL) and PEO-TSA (10 mL) solutions were loaded into individual syringes, and the two solutions were electrospun simultaneously onto a centrally located mandrel. Scaffolds with fiber fractions of 50-55% PEO (or PEO-TSA) fibers were used for experiments.

### Immunofluorescent Imaging of Histone Acetylation

After 24 hours of culture on tissue culture plastic in basal media, pMFCs were treated with or without medium-dose of 325 nM TSA for 24 hours. Cells were fixed in 1:1 solution of ice cold ethanol/methanol followed by washing with PBS. Fixed cells were blocked 5% goat serum for 24 hours, immunostained with an acetyl-Histone H3 (Lys9) rabbit antibody (Ac-H3K9, MA5-11195, 1:400, Thermo Fisher Scientific Inc., Waltham, MA) for 24 hours, and fluorescently labeled with goat anti-rabbit, Alexa Fluor-488 secondary antibody (A-11008, Invitrogen). Cells were counterstained with 4’, 6-diamidno-2-phenylindole (DAPI) and imaged on an inverted microscope. The mean fluorescence intensity per cell was measured (ImageJ)^42^.

### Chromatin condensation parameter and nuclear deformability

pMFCs were seeded on aligned nanofibrous PCL scaffolds and cultured for 2 days in basal media. To induce chromatin decondensation, cells were treated with medium-dose TSA for 24 hours. To quantify the chromatin condensation parameter (CCP), cell-laden scaffolds were fixed with paraformaldehyde followed by washing with PBS and subsequent permeabilization with 0.05% Triton X-100. Nuclei were labeled with DAPI (ProLong® Gold with DAPI, P36935, Molecular Probes®, Grand Island, NY) and imaged at their mid-section using confocal. Sobel edge detection (MATLAB) was used to calculate the edge density within individual nuclei and the resulting CCP^13,32,43,44^.

Nuclear deformability was quantified using live cells on scaffolds. To assess nuclear deformability, the nuclear aspect ratio (NAR) was calculated after applying 0, 3, 6, 9, 12, and 15% grip-to-grip static strain to the cell-seeded scaffolds using a custom device^45,13^. At each strain level, DAPI-stained nuclei were imaged on an inverted microscope (Nikon T30, Nikon Instruments, Melville, NY). NAR was calculated using a custom algorithm (MATLAB)^13^, and changes in NAR were tracked for individual nuclei at each strain level.

### In vitro cell migration

To query 2D pMFC migration following chromatin decondensation, a standard 2-dimensional monolayer scratch assay was performed with and without TSA treatment. For scratch assays, pMFCs were seeded at high density (20,000 cells/cm^2^) on 6-well tissue culture plastic and cultured to confluency for 2 days prior to chromatin decondensation with medium-dose TSA for 24 hours. Confluent monolayers with or without TSA treatment were then scratched using a 200 µL pipette tip, followed by washing with PBS to remove cell debris. Using an inverted microscope, the scratch areas were imaged at time 0 and every 4 hours until scratch closure, and the scratch areas were computed using ImageJ^42^.

### Cell migration into devitalized meniscus ECM

To assess 3-dimensional migration of endogenous pMFCs into devitalized native meniscus extracellular matrix (ECM), cylindrical meniscus explants (4 mm diameter, 2 mm height) were biopsied from freshly dissected porcine menisci. Explants were incubated in basal media for 2 weeks with media changes every other day. To obtain devitalized native meniscus tissue ECM cryosections, whole menisci were cryopreserved and embedded in Optimal Cutting Temperature sectioning medium (OCT; Sakura Finetek, Torrance, CA). The OCT-embedded whole menisci were cryosectioned axially (i.e., longitudinal cryosections) at 40 µm thickness, mounted on positively charged glass slides, and stored at -20ºC.

After 2 weeks of in vitro culture, the cells within explants were fluorescently labeled with 5 µg/mL of 5-chloromethylfluorescein diacetate (CellTracker™ Green; Thermo Fisher Scientific Inc., Waltham, MA) in serum-free media (DMEM with 1% PSF) for 1 hour. Subsequently, the fluorescently labeled explants were incubated in serum-free media for 30 min and then rinsed with PBS. Devitalized meniscus ECM cryosections were thawed at room temperature, washed in PBS, and sterilized under ultraviolet light for 1 hour. Explants were set on top of the sterile devitalized meniscus ECM cryosections and incubated at 37°C with or without medium-dose TSA treatment in basal media for 2 days. At the 2 day end point, explants were removed, and maximum z-stack projections of the devitalized tissue substrate were acquired using a confocal microscope. Using a custom algorithm (MATLAB), the cell infiltration depth was measured as the distance from the explant-ECM interface maximum cell within the devitalized meniscus ECM^4,10,11,13^. The total number of cells and the number of migrated cells (those entirely embedded within the devitalized meniscus ECM) were counted using ImageJ^42^.

### RNA-sequencing

Aligned nanofibrous PCL scaffolds (1.5 × 1 cm and ∼500 µm thick) were hydrated in PBS and coated in fibronectin (20 µg/mL) for ≥24 hours. pMFCs from 3 porcine donors (N=3) were isolated as above and seeded onto scaffolds for 24 hours prior to treatment. Scaffolds were divided into 6 groups based on treatment regimen: (Group 1) Vehicle control at <0.1 % dimethyl sulfoxide (DMSO), (Groups 2 & 3) TSA (low-dose: 150 nM & high-dose: 650 nM), (Group 4) TGFβ3 at 10 ng/mL, and (Groups 5 & 6) TSA + TGFβ3 (with the same doses as groups 2 and 3). Low (150 nM) and high (650nM) TSA concentrations were determined based on prior work^13^, which showed that 150nM was the lowest TSA dose to increase meniscus cell migration through porous membranes compared to DMSO-treated controls.

DMSO or TSA was delivered for 24 hours and then washed out with extended incubation in basal media up to day 7 of culture. On day 4 of culture (i.e., 3 days of culture in basal media after TSA treatment), TGFβ3 was delivered to stimulate matrix production (Groups 4, 5, and 6). Study endpoints included 24 hours after seeding (i.e., time 0 before treatment for group 1); at 1 day immediately following DMSO or TSA treatment for groups 1, 2, and 3; and at 7 days after treatment for all groups. At each study endpoint, RNA was isolated following the manufacturer’s protocol (mRNeasy Plus Mini Kit, Qiagen). RNA concentrations were quantified (Nanodrop) and subsequently analyzed via fragment analysis (BioAnalyzer, Agilent, all RIN > 8). RNA libraries were prepared using Illumina polyA total RNA kit. Single-end reads were sequenced using Illumina NovaSeq (100 base pairs), and reads were aligned to the sscrofa11.1 genome via HiSat2^46,47^. Picard and FeatureCounts were used to generate bam files and count files, respectively^48^. For differential gene expression analysis between groups, Deseq2 was used in R Studio, and Venn diagrams were generated^49^. For gene ontology, Panther and the Gene Ontology Resource were used^33–35^.

### Nascent matrix labeling

The same groups used for RNA-sequencing were used for nascent matrix labeling. To stain the nascent matrix produced by seeded cells from day 4 to study endpoints at day 7 or 14, cell-seeded scaffolds were cultured in defined media including the noncanonical amino acid azidohomoalanine (AHA) and methionine at a ratio 3:1, respectively. During the culture duration, the cells incorporate the azide-modified noncanonical methionine analog into cell-produced matrix (i.e., the nascent matrix). This enables subsequent click labeling with azide-dibenzo cyclooctyne (DBCO)^38–41^. At each end point, cell-laden scaffolds were fixed with paraformaldehyde followed by washing with phosphate buffered saline (PBS). Subsequently, newly formed nascent matrix, nuclei, and actin were fluorescently labeled via incubation with Alexa Fluor (AF)488-DBCO, Draq5, and Phalloidin-AF546, respectively. Cell-produced nascent matrix, nuclei, actin, and scaffolds (auto-fluorescence) were imaged via confocal.

### TSA delivery via dual material scaffold

In order to develop a rapid and local delivery mechanism for TSA within the injury site, we fabricated a dual material nanofibrous scaffold with PCL and PEO as described above. The water soluble PEO component was loaded with TSA for rapid release upon implantation. To test the effectiveness of scaffold-delivered TSA on endogenous cells, we employed an ex vivo meniscus injury model.

Whole porcine menisci were subjected to a vertical defect approximately 1 cm in length. PCL/PEO scaffolds with and without TSA (PCL/PEO and PCL/PEO-TSA scaffolds, N=5/group) were secured within the defects via a vertical mattress suture and incubated in basal media for 1 or 7 days. At each timepoint, constructs were embedded in OCT and cryosectioned perpendicular to the defect (i.e., radial cryosections) to 10 µm thickness, mounted on positively charged glass slides, and stored at -20 ºC.

For quantification of cell acetylation and migration, cryosections were thawed, washed in PBS, and fixed in paraformaldehyde. For cell acetylation, fixed sections were permeabilized with 0.1% triton X at room temperature. After blocking in 10% bovine serum albumin for 30 min at room temperature, sections were immunostained for acetyl-Histone H3 (Lys9) rabbit antibody (Ac-H3K9, MA5-11195, 1:400, Thermo Fisher Scientific Inc., Waltham, MA) and fluorescently labeled with goat anti-rabbit, Alexa Fluor-488 secondary antibody (Catalog #: A-11008, Invitrogen) and then Ac-H3K9 immunofluorescence was imaged via confocal. For quantification of cell migration, fixed sections were stained Hematoxylin and Eosin and then imaged at the peripheral and central wound edges. Total cell number was calculated using a custom MATLAB script^50,51^.

### Statistical analyses

Statistical analyses were performed using t-tests for comparisons of control versus TSA or one-way ANOVA for normalized data across time points with Tukey’s post hoc test (Prism 9, Graphpad). Data are represented as the mean ± the standard deviation unless otherwise noted, and differences were considered statistically significant at p-value < 0.05. For differential expression analysis, genes were considered differentially expressed if both the adjusted p-value < 0.05 and an absolute fold change in expression between groups > 2. The fold change for matched differentially expressed genes between two groups were fit to a simple linear regression and Pearson’s coefficient was calculated.

### Data availability

The generated raw and processed data from the RNA-seq experiments will be available in the Gene Expression Omnibus (GEO) repository or by request from the corresponding author. All other data are available upon request from the corresponding authors.

## Acknowledgements

The National Institutes of Health (R01 AR056624, R01 AR071340, L30HL165571, and T32 AR053461), the Penn Center for Musculoskeletal Disorders (P30 AR069619), and the Department of Veterans Affairs (I01 RX003375 and IK1 RX003932) supported this work. The authors thank the Penn Genomic Sequencing Core for technical assistance with RNA-sequencing. We gratefully thank the members of the Mauck, Gullbrand, and Heo labs for their thoughtful discussions and feedback during conceptualization and data analysis.

## Author contributions

The contributions for each author are as described. *Ryan Locke* and *Liane Miller*: Both authors contributed equally to conceptualization, methodology, investigation, analysis, validation, writing – original draft, writing – review and editing, visualization. *Beth Lemmon, Sereen Assi, Dakota Jones, Eddie Bonnevie*: Methodology, investigation, analysis, writing – original draft, writing – review and editing, visualization. *Jason Burdick*: Conceptualization, methodology, funding acquisition, writing – review and editing. *Su Jin Heo and Robert Mauck*: Conceptualization, methodology, funding acquisition, resources, writing – original draft, writing – review and editing.

## Supplemental data

**Supplemental Figure 1.**
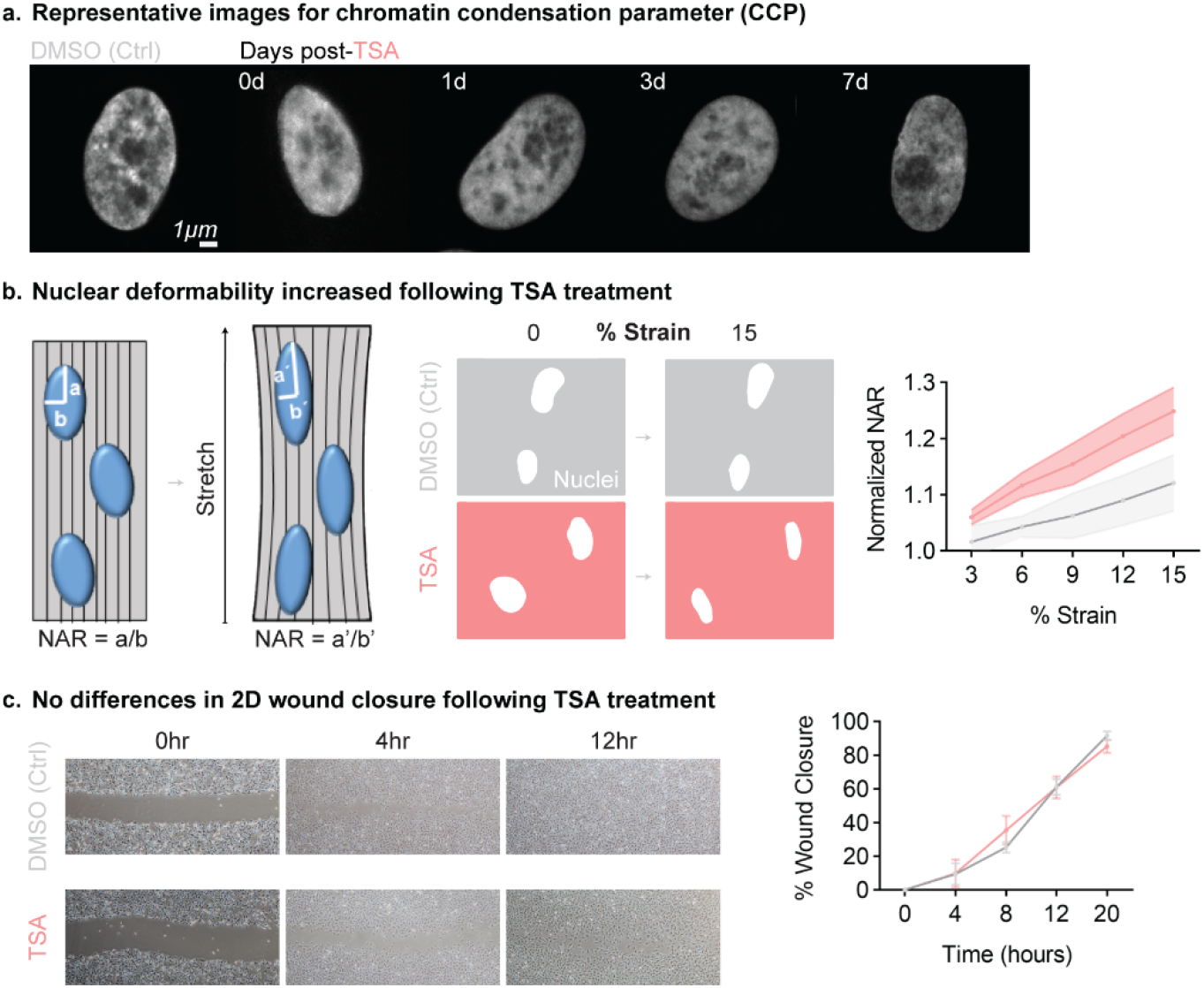
(a) Representative images of high magnification DAPI-stained nuclei used for quantification of chromatin condensation parameter edge density. (b) Schematic of nuclear deformability assay and representative binarized images of DAPI-stained nuclei at 0 and 15% strain for DMSO-treated controls and TSA-treated cells. Quantification of the nuclear aspect ratio (NAR) at incremental strain steps for DMSO-treated controls and TSA-treated cells, showing increased nuclear deformability at each strain step following TSA treatment. (c) Representative images of 2-dimensional scratch assay over time for DMSO-treated controls and TSA-treated cells, and quantification of percent wound closure over time. N=5-6.

**Supplemental Figure 2.**
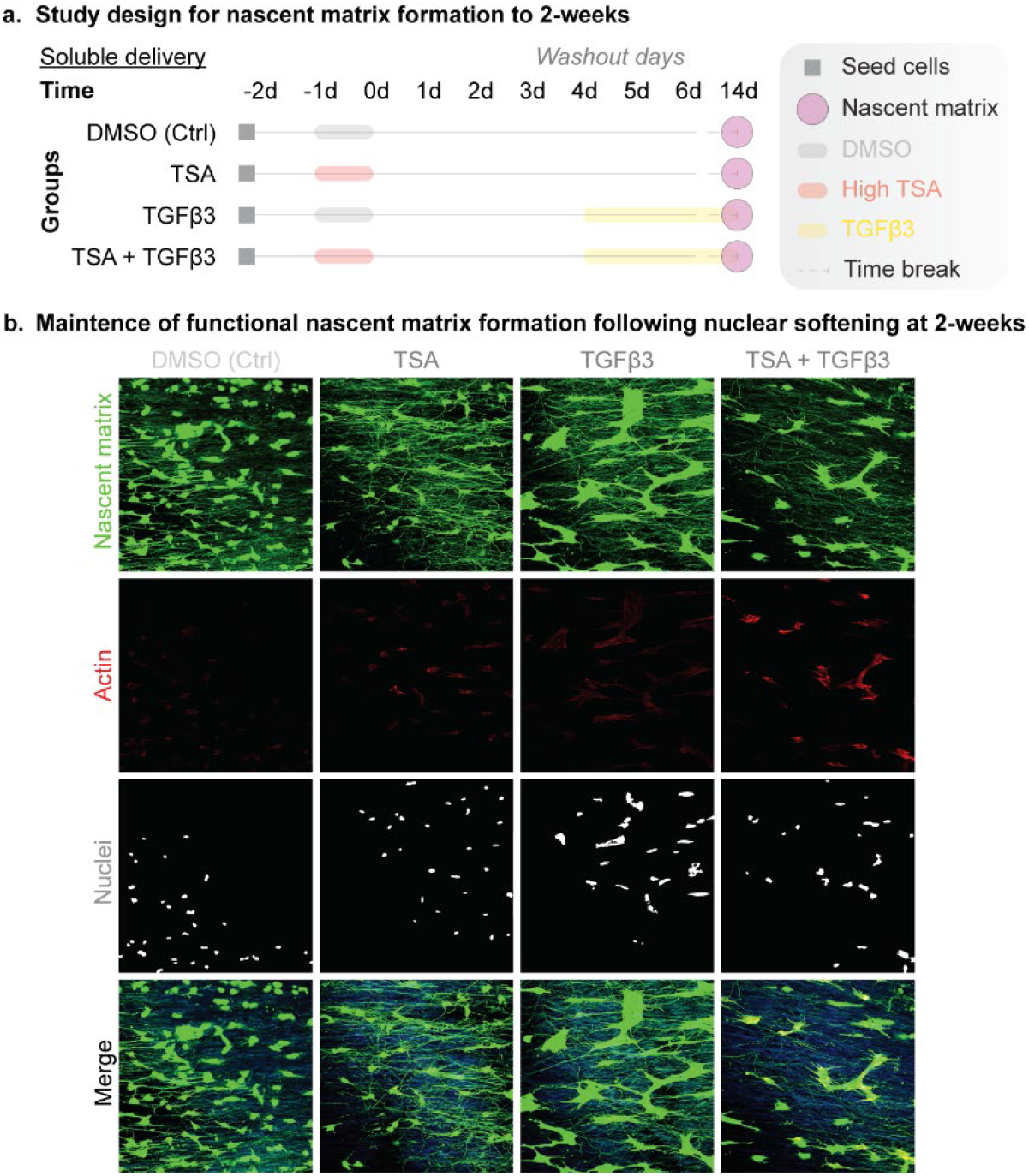
Representative confocal images of nascent matrix at 14 days following DMSO, TSA, TGFβ3, or TSA + TGFβ3 treatment. Green: deposited nascent matrix over the culture period, yellow: overlay of nascent matrix and actin (red), blue: nanofibrous scaffold autofluorescence, magenta: nuclei.

**Supplemental Figure 3.**
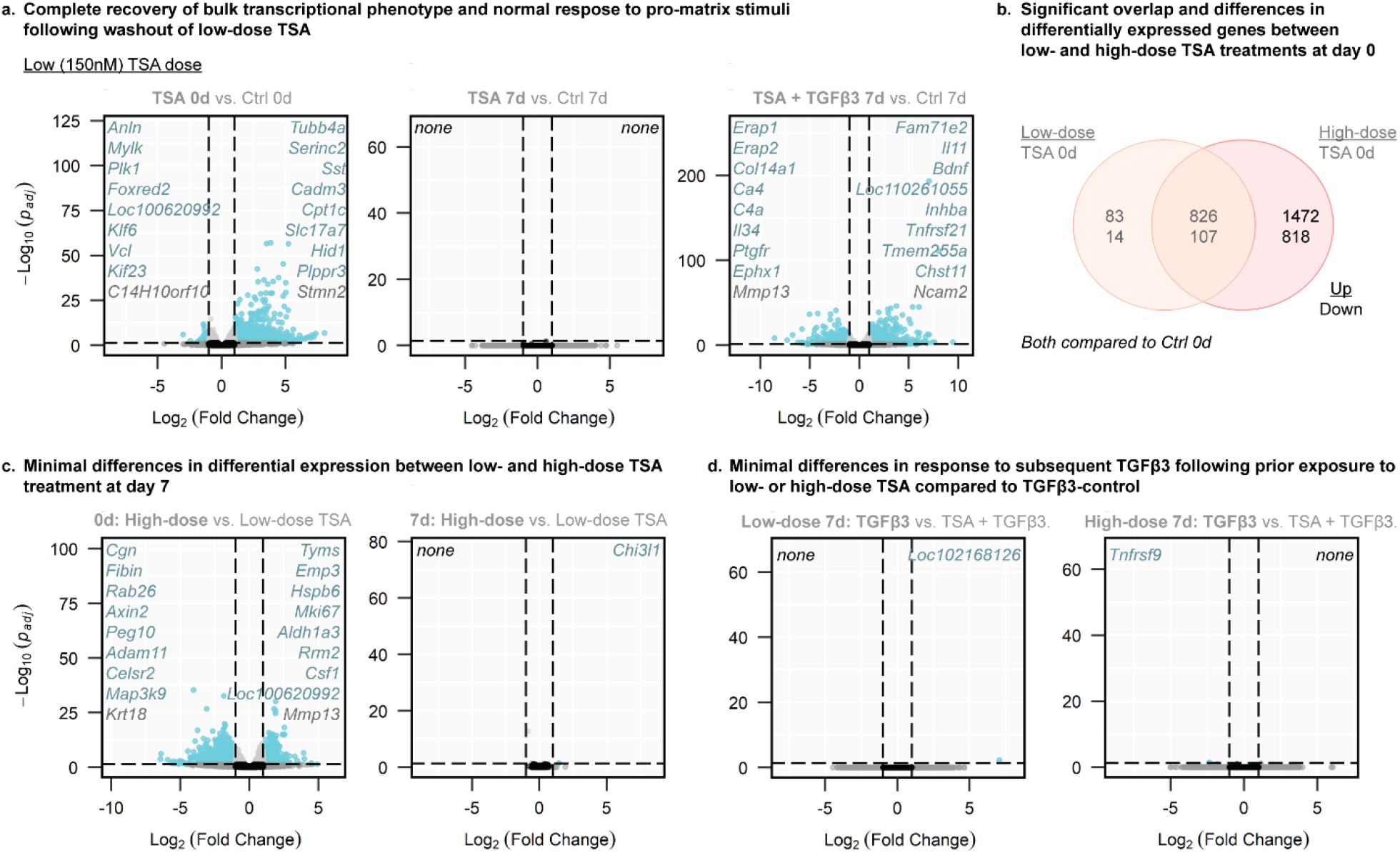
(a) Volcano plots for low-dose TSA treatment groups. (b) Venn diagram comparing differentially expressed genes at day 0 following low- and high-dose TSA. Numbers indicate upregulated (top number) genes and downregulated (bottom number) genes. Non-overlapping quadrants represent unique genet sets for each group. (c) Volcano plots for high-dose compared to low-dose TSA treatment at day 0 and 7. (d) Volcano plots for TGFβ3 only compared to low- or high-dose TSA + TGFβ3 at day 7.

**Supplemental Figure 4.**
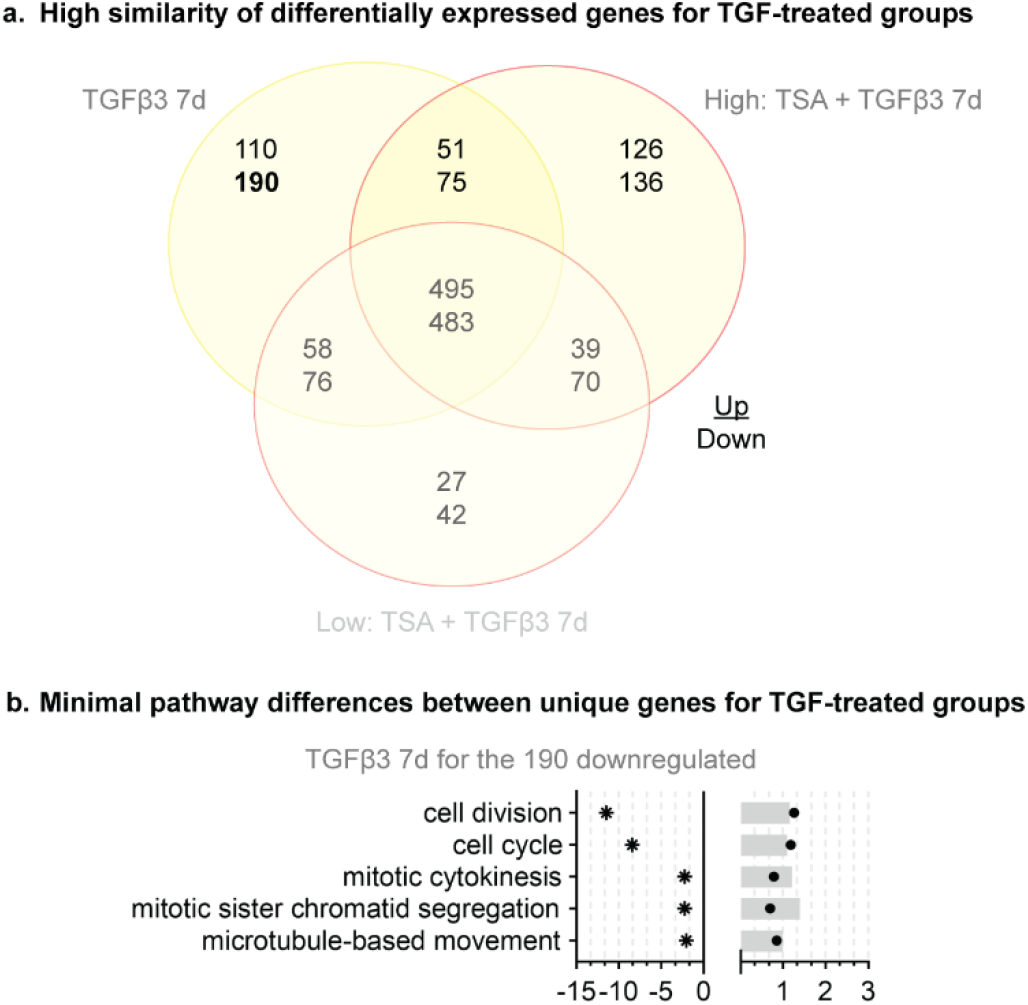
(a) Venn diagram of differentially expressed genes between groups treated with TGFβ3, including TGFβ3 only, high-dose TSA + TGFβ3, and low-dose TSA + TGFβ3. Numbers indicate upregulated (top number) genes and downregulated (bottom number) genes. Non-overlapping quadrants represent unique gene sets for each group. (b) Gene ontologies for unique upregulated genes following high-dose TSA + TGFβ3. Note: The unique downregulated genes following high-dose TSA + TGFβ3 and both the up- and down-regulated genes following low-dose TSA + TGFβ3 were not significantly enriched into specific gene ontologies.

## References

1. Gurtner, G. C., Werner, S., Barrandon, Y. & Longaker, M. T. Wound repair and regeneration. Nature 453, 314–321 (2008).

2. Longo, S. K., Guo, M. G., Ji, A. L. & Khavari, P. A. Integrating single-cell and spatial transcriptomics to elucidate intercellular tissue dynamics. Nat. Rev. Genet. 22, 627–644 (2021).

3. Qu, F., Guilak, F. & Mauck, R. L. Cell migration: implications for repair and regeneration in joint disease. Nat. Rev. Rheumatol. 15, 167–179 (2019).

4. Qu, F. et al. Maturation State and Matrix Microstructure Regulate Interstitial Cell Migration in Dense Connective Tissues. Sci. Rep. 8, 3295 (2018).

5. McGregor, A. L., Hsia, C.-R. & Lammerding, J. Squish and squeeze—the nucleus as a physical barrier during migration in confined environments. Curr. Opin. Cell Biol. 40, 32–40 (2016).

6. Bansal, S. et al. Meniscal repair: The current state and recent advances in augmentation. J. Orthop. Res. 39, 1368–1382 (2021).

7. Makris, E. A., Hadidi, P. & Athanasiou, K. A. The knee meniscus: Structure–function, pathophysiology, current repair techniques, and prospects for regeneration. Biomaterials 32, 7411–7431 (2011).

8. Makris, E. A., Gomoll, A. H., Malizos, K. N., Hu, J. C. & Athanasiou, K. A. Repair and tissue engineering techniques for articular cartilage. Nat. Rev. Rheumatol. 11, 21–34 (2015).

9. Rai, M. F. & Sandell, L. J. Regeneration of articular cartilage in healer and non-healer mice. Matrix Biol. J. Int. Soc. Matrix Biol. 39, 50–55 (2014).

10. Qu, F., Holloway, J. L., Esterhai, J. L., Burdick, J. A. & Mauck, R. L. Programmed biomolecule delivery to enable and direct cell migration for connective tissue repair. Nat. Commun. 8, 1780 (2017).

11. Qu, F. et al. Repair of dense connective tissues via biomaterial-mediated matrix reprogramming of the wound interface. Biomaterials 39, 85–94 (2015).

12. Martin, A. R. et al. Nanofibrous hyaluronic acid scaffolds delivering TGF-β3 and SDF-1α for articular cartilage repair in a large animal model. Acta Biomater. 126, 170–182 (2021).

13. Heo, S.-J. et al. Nuclear softening expedites interstitial cell migration in fibrous networks and dense connective tissues. Sci. Adv. 6, eaax5083 (2020).

14. Travers, J. G. et al. HDAC Inhibition Reverses Preexisting Diastolic Dysfunction and Blocks Covert Extracellular Matrix Remodeling. Circulation 143, 1874–1890 (2021).

15. Das, A., Barai, A., Monteiro, M., Kumar, S. & Sen, S. Nuclear softening is essential for protease-independent migration. Matrix Biol. 82, 4–19 (2019).

16. Nava, M. M. et al. Heterochromatin-Driven Nuclear Softening Protects the Genome against Mechanical Stress-Induced Damage. Cell 181, 800-817.e22 (2020).

17. Mukherjee, A., Barai, A., Singh, R. K., Yan, W. & Sen, S. Nuclear plasticity increases susceptibility to damage during confined migration. PLOS Comput. Biol. 16, e1008300 (2020).

18. dos Santos, Á. et al. DNA damage alters nuclear mechanics through chromatin reorganization. Nucleic Acids Res. 49, 340–353 (2021).

19. Walker, C. J. et al. Nuclear mechanosensing drives chromatin remodelling in persistently activated fibroblasts. Nat. Biomed. Eng. 5, 1485–1499 (2021).

20. Alisafaei, F., Jokhun, D. S., Shivashankar, G. V. & Shenoy, V. B. Regulation of nuclear architecture, mechanics, and nucleocytoplasmic shuttling of epigenetic factors by cell geometric constraints. Proc. Natl. Acad. Sci. 116, 13200–13209 (2019).

21. Chalut, K. J. et al. Chromatin decondensation and nuclear softening accompany Nanog downregulation in embryonic stem cells. Biophys. J. 103, 2060–2070 (2012).

22. Krause, M. et al. Cell migration through three-dimensional confining pores: speed accelerations by deformation and recoil of the nucleus. Philos. Trans. R. Soc. Lond. B. Biol. Sci. 374, 20180225 (2019).

23. Friedl, P., Sahai, E., Weiss, S. & Yamada, K. M. New dimensions in cell migration. Nat. Rev. Mol. Cell Biol. 13, 743–747 (2012).

24. Guilak, F., Tedrow, J. R. & Burgkart, R. Viscoelastic Properties of the Cell Nucleus. Biochem. Biophys. Res. Commun. 269, 781–786 (2000).

25. Lautscham, L. A. et al. Migration in Confined 3D Environments Is Determined by a Combination of Adhesiveness, Nuclear Volume, Contractility, and Cell Stiffness. Biophys. J. 109, 900–913 (2015).

26. van Helvert, S., Storm, C. & Friedl, P. Mechanoreciprocity in cell migration. Nat. Cell Biol. 20, 8–20 (2018).

27. Calero-Cuenca, F. J., Janota, C. S. & Gomes, E. R. Dealing with the nucleus during cell migration. Curr. Opin. Cell Biol. 50, 35–41 (2018).

28. Friedl, P., Wolf, K. & Lammerding, J. Nuclear mechanics during cell migration. Curr. Opin. Cell Biol. 23, 55–64 (2011).

29. Gerlitz, G. & Bustin, M. The role of chromatin structure in cell migration. Trends Cell Biol. 21, 6–11 (2011).

30. Srivatsan, S. R. et al. Massively multiplex chemical transcriptomics at single-cell resolution. Science 367, 45–51 (2020).

31. Gurgul, A. et al. The effect of histone deacetylase inhibitor trichostatin A on porcine mesenchymal stem cell transcriptome. Biochimie 139, 56–73 (2017).

32. Irianto, J., Lee, D. A. & Knight, M. M. Quantification of chromatin condensation level by image processing. Med. Eng. Phys. 36, 412–417 (2014).

33. Mi, H., Muruganujan, A., Ebert, D., Huang, X. & Thomas, P. D. PANTHER version 14: more genomes, a new PANTHER GO-slim and improvements in enrichment analysis tools. Nucleic Acids Res. 47, D419–D426 (2019).

34. Ashburner, M. et al. Gene Ontology: tool for the unification of biology. Nat. Genet. 25, 25–29 (2000).

35. Gene Ontology Consortium. The Gene Ontology resource: enriching a GOld mine. Nucleic Acids Res. 49, D325–D334 (2021).

36. Huang, D. W., Sherman, B. T. & Lempicki, R. A. Systematic and integrative analysis of large gene lists using DAVID bioinformatics resources. Nat. Protoc. 4, 44–57 (2009).

37. Sherman, B. T. et al. DAVID: a web server for functional enrichment analysis and functional annotation of gene lists (2021 update). Nucleic Acids Res. 50, W216–W221 (2022).

38. Loebel, C., Mauck, R. L. & Burdick, J. A. Local nascent protein deposition and remodelling guide mesenchymal stromal cell mechanosensing and fate in three-dimensional hydrogels. Nat. Mater. 18, 883–891 (2019).

39. Loebel, C. et al. Metabolic Labeling to Probe the Spatiotemporal Accumulation of Matrix at the Chondrocyte–Hydrogel Interface. Adv. Funct. Mater. 30, 1909802 (2020).

40. Loebel, C. et al. Metabolic labeling of secreted matrix to investigate cell–material interactions in tissue engineering and mechanobiology. Nat. Protoc. 17, 618–648 (2022).

41. Patel, J. M. et al. Stabilization of Damaged Articular Cartilage with Hydrogel-Mediated Reinforcement and Sealing. Adv. Healthc. Mater. 10, 2100315 (2021).

42. Schindelin, J. et al. Fiji: an open-source platform for biological-image analysis. Nat. Methods 9, 676–682 (2012).

43. Heo, S.-J. et al. Differentiation alters stem cell nuclear architecture, mechanics, and mechano-sensitivity. eLife 5, e18207 (2016).

44. Heo, S.-J. et al. Aberrant chromatin reorganization in cells from diseased fibrous connective tissue in response to altered chemomechanical cues. Nat. Biomed. Eng. 1–15 (2022) doi:10.1038/s41551-022-00910-5.

45. Bonnevie, E. D. et al. Aberrant mechanosensing in injured intervertebral discs as a result of boundary-constraint disruption and residual-strain loss. Nat. Biomed. Eng. 3, 998–1008 (2019).

46. Warr, A. et al. An improved pig reference genome sequence to enable pig genetics and genomics research. GigaScience 9, giaa051 (2020).

47. Kim, D., Paggi, J. M., Park, C., Bennett, C. & Salzberg, S. L. Graph-based genome alignment and genotyping with HISAT2 and HISAT-genotype. Nat. Biotechnol. 37, 907–915 (2019).

48. Liao, Y., Smyth, G. K. & Shi, W. featureCounts: an efficient general purpose program for assigning sequence reads to genomic features. Bioinformatics 30, 923–930 (2014).

49. Love, M. I., Huber, W. & Anders, S. Moderated estimation of fold change and dispersion for RNA-seq data with DESeq2. Genome Biol. 15, 550 (2014).

50. David, M. A. et al. Early, focal changes in cartilage cellularity and structure following surgically induced meniscal destabilization in the mouse. J. Orthop. Res. 35, 537–547 (2017).

51. Lemmon, E. A., Locke, R. C., Szostek, A. K., Ganji, E. & Killian, M. L. Partial-width injuries of the rat rotator cuff heal with fibrosis. Connect. Tissue Res. 59, 437–446 (2018).

