## Supplemental data for "Rapid Restoration of Cell Phenotype and Matrix Forming Capacity Following Transient Nuclear Softening"

### Title


### Supplemental data

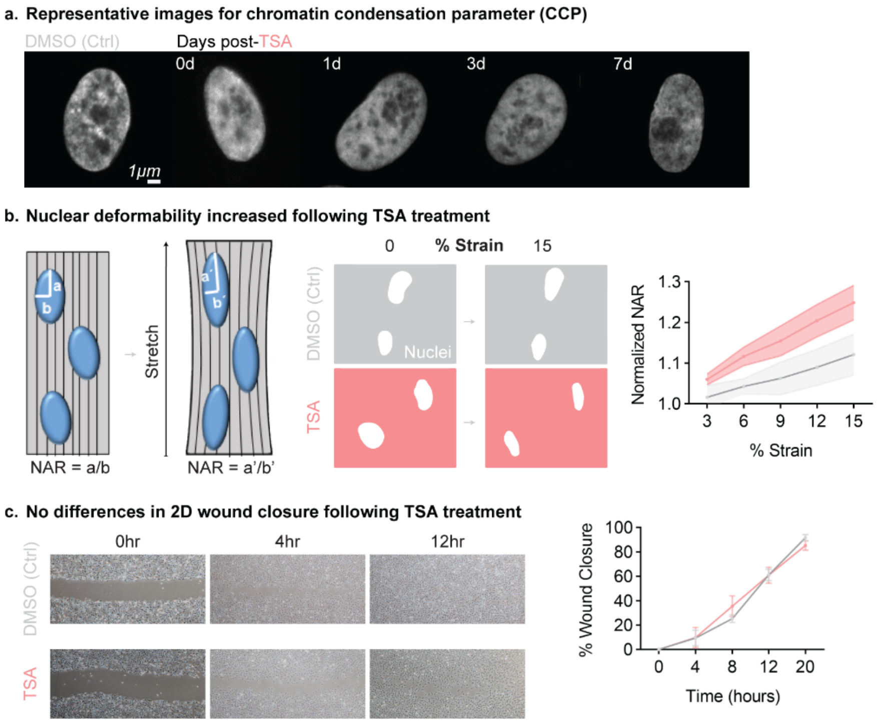


**Supplemental Figure 1.** (a) Representative images of high magnification DAPI-stained nuclei used for quantification of chromatin condensation parameter edge density. (b) Schematic of nuclear deformability assay and representative binarized images of DAPI-stained nuclei at 0 and 15% strain for DMSO-treated controls and TSA-treated cells. Quantification of the nuclear aspect ratio (NAR) at incremental strain steps for DMSO-treated controls and TSA-treated cells, showing increased nuclear deformability at each strain step following TSA treatment. (c) Representative images of 2-dimensional scratch assay over time for DMSO-treated controls and TSA-treated cells, and quantification of percent wound closure over time. N=5-6.

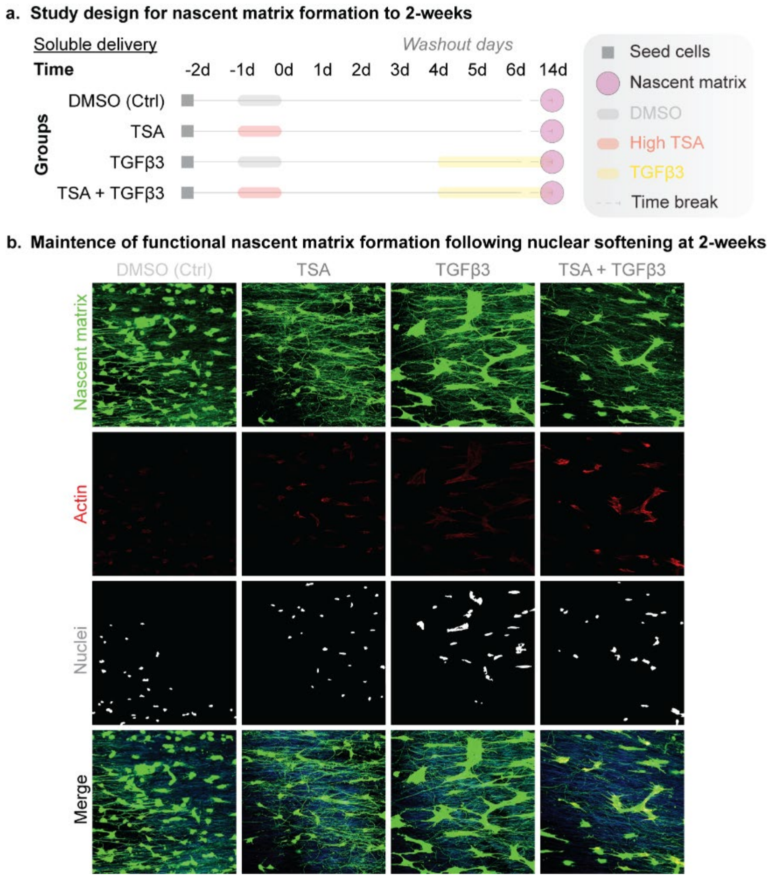


**Supplemental Figure 2.** Representative confocal images of nascent matrix at 14 days following DMSO, TSA, TGFβ3, or TSA + TGFβ3 treatment. Green: deposited nascent matrix over the culture period, yellow: overlay of nascent matrix and actin (red), blue: nanofibrous scaffold autofluorescence, magenta: nuclei.

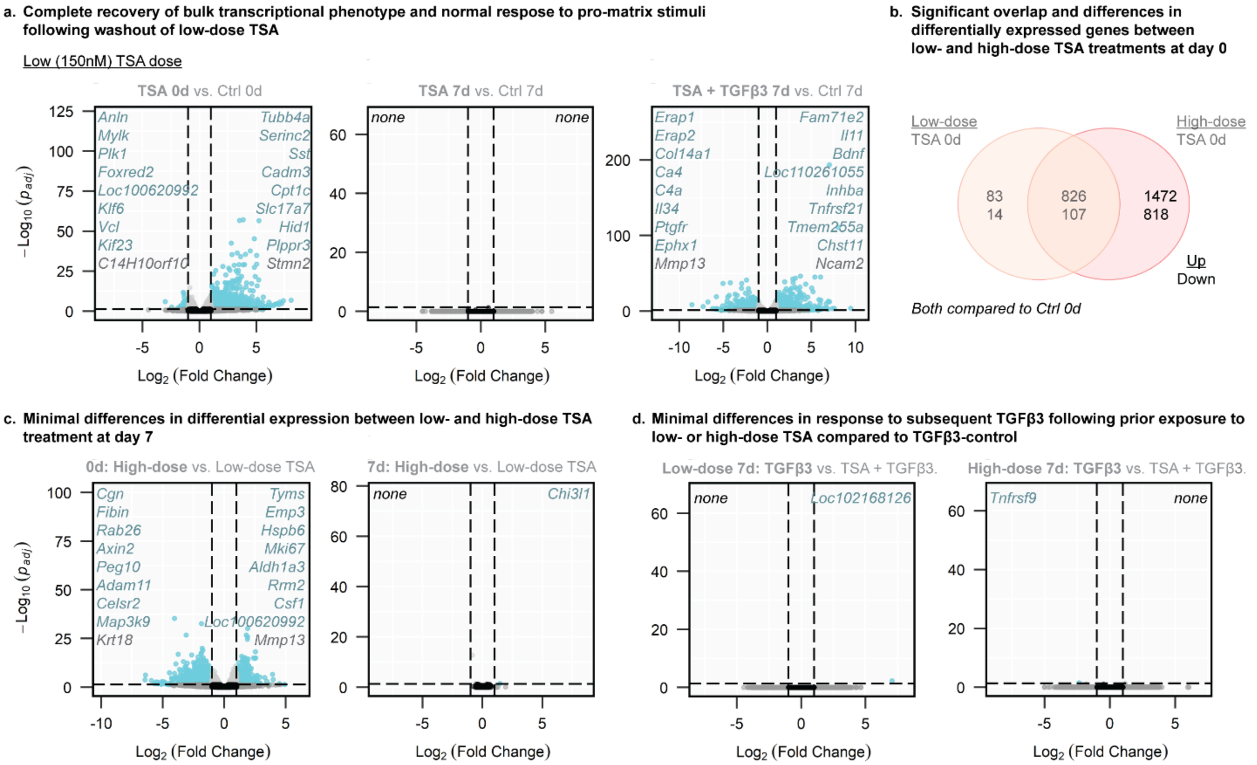


**Supplemental Figure 3.** (a) Volcano plots for low-dose TSA treatment groups. (b) Venn diagram comparing differentially expressed genes at day 0 following low- and high-dose TSA. Numbers indicate upregulated (top number) genes and downregulated (bottom number) genes. Non-overlapping quadrants represent unique genet sets for each group. (c) Volcano plots for high-dose compared to low-dose TSA treatment at day 0 and 7. (d) Volcano plots for TGFβ3 only compared to low- or high-dose TSA + TGFβ3 at day 7.

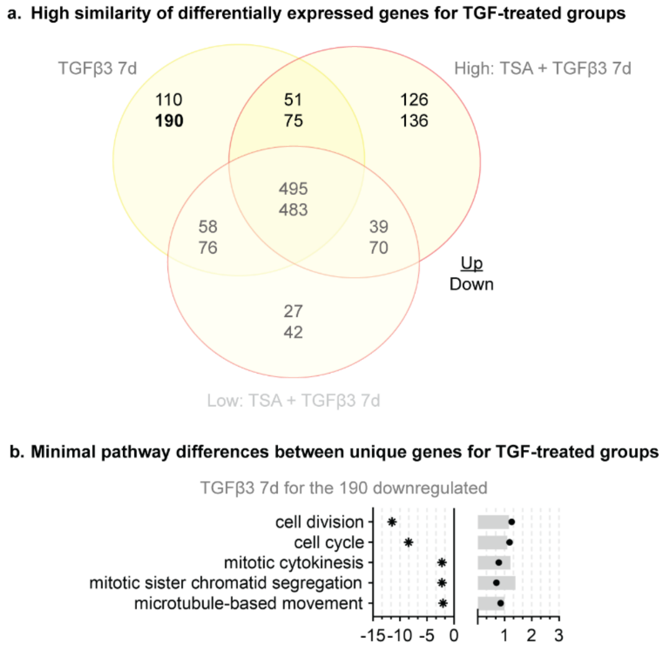


**Supplemental Figure 4.** (a) Venn diagram of differentially expressed genes between groups treated with TGFβ3, including TGFβ3 only, high-dose TSA + TGFβ3, and low-dose TSA + TGFβ3. Numbers indicate upregulated (top number) genes and downregulated (bottom number) genes. Non-overlapping quadrants represent unique gene sets for each group. (b) Gene ontologies for unique upregulated genes following high-dose TSA + TGFβ3. Note: The unique downregulated genes following high-dose TSA + TGFβ3 and both the up- and down-regulated genes following low-dose TSA + TGFβ3 were not significantly enriched into specific gene ontologies.
